# Pre-stimulus Brain States Predict and Control Variability in Stimulation Responses

**DOI:** 10.1101/2025.09.18.677069

**Authors:** Giovanni Rabuffo, Marianna Angiolelli, Tomoki Fukai, Gustavo Deco, Pierpaolo Sorrentino, Davide Momi

## Abstract

Does the ongoing brain state determine how it responds to localized electrical stimulation? This fundamental question has major implications for neuroscience and medicine, as stimulation outcomes remain highly variable even when identical parameters (*how*) and target sites (*where*) are used. Such unpredictability undermines reproducibility, limits clinical reliability, and forces current protocols to rely on empirical trial-and-error rather than principled, evidence-based strategies. Mounting evidence suggests that the brain’s state prior to stimulation–a crucial *when* factor–shapes responses, yet the most reliable predictive markers remain unknown. Here, we systematically characterize the links between pre-stimulus (spontaneous) activity and post-stimulus (evoked) responses using simultaneous high-density EEG and stereotactic EEG from 36 epilepsy patients across *∼*320 sessions (*>*10,000 individual stimulations). We show that large-scale neural dynamics robustly predict stimulation outcomes, with a subset of measures–particularly network synchronization, functional connectivity, and spatiotemporal signal diversity–consistently forecasting responses across sessions. Whole-brain activity enhanced prediction compared to local assessments, and predictability varied across networks, being strongest in sensorimotor and visual regions. These findings establish a quantitative framework for state-dependent brain stimulation: by timing interventions to optimal pre-stimulus states, variability can be reduced and reproducibility enhanced. Our results directly address the fundamental question of *where* and *when* to stimulate, providing a pathway toward evidence-based protocols with improved therapeutic precision.

## 1 Introduction

Brain stimulation techniques offer powerful tools for both investigating neural mechanisms and treating neurological disorders. Yet, their clinical potential remains limited by a fundamental problem: stimulation effects vary unpredictably across trials, even with identical parameters [1–5]. This variability undermines diagnostic precision and therapeutic reliability, forcing clinicians to rely on empirical approaches rather than principled, evidence-based protocols [6]. Understanding stimulation variability requires considering three key factors: where to stimulate (anatomical target), how to stimulate (stimulus parameters), and, crucially, when to stimulate (brain state timing) [7, 8]. Although advances in neuroimaging have refined target selection and stimulus optimization has improved parameter choices [9], the temporal dimension remains poorly understood. Even when anatomical targets and stimulus parameters are identical, responses can vary by several-fold across trials [1, 10–12], suggesting that the brain’s ongoing state at the moment of stimulation plays a decisive role in determining outcomes.

Recent evidence demonstrates that brain state before stimulation systematically influences evoked responses [13–15]. For instance, the phase of ongoing oscillations in target regions can double or halve evoked potential amplitudes (e.g., [16–18]). Similarly, pre-stimulus connectivity patterns can be targeted to modulate post-stimulus effects [19]. However, current closed-loop systems rely on simple markers, typically oscillatory phase in single regions, while the optimal features for predicting stimulation outcomes remain unknown.

The most widely used brain stimulation methods are applied without accounting for the dynamic nature of pre-stimulus (spontaneous) brain activities, leading to significant variability in post-stimulus (evoked) responses and hindering the efficacy of clinical applications. However, the spontaneous dynamics are fine-tuned in the healthy individuals [20], and affected by neurological diseases [21–23], which suggests that they are behaviorally relevant and should not be neglected. In other words, the relationship between spontaneous and evoked activity likely reflects fundamental principles of brain organization, relevant for therapeutic design.

We hypothesize that pre-stimulus network states constrain post-stimulus responses through competitive resource allocation: when neural populations maintain strong functional connections, fewer resources remain available for stimulus-driven reorganization [24]. From a dynamical systems perspective, ongoing activity may reflect metastable attractor states that resist perturbation unless external input exceeds critical thresholds [25–27]. This framework is supported by evidence showing that large-scale interactions across regions are best understood in terms of the spreading of high-amplitude bursts [28, 29]. Cortical regions exhibit distinct excitability gradients and recurrent connectivity patterns [30], which may differentially influence their susceptibility to stimulation-induced perturbations. This framework predicts that specific spontaneous configurations should be more amenable to stimulation-induced changes than others, with neuronal populations already engaged in coherent functional states exhibiting reduced sensitivity to incoming stimuli.

To answer this fundamental question systematically, we quantified trial-by-trial relationships between pre- and post-stimulus brain dynamics. Unlike traditional approaches that suppress variability through trial-averaging, we treat spontaneous fluctuations as meaningful signals containing predictive information about individual responses to brain stimulation. We employed a comprehensive analytical approach spanning statistical measures (capturing local signal dynamics), network measures (quantifying large-scale coordination), and information-theoretic measures (evaluating signal complexity and predictability). This multi-domain strategy was designed to capture complementary aspects of neural organization that no single framework could reveal alone.

Using simultaneous intracranial and high-density EEG recordings from 36 patients undergoing clinical stimulation [31, 32], we systematically tested three key predictions: (1) spontaneous brain states reliably predict evoked responses across trials, (2) whole-brain context provides stronger prediction than local measurements alone, and (3) predictability changes according to the brain network being stimulated. Our findings establish a quantitative foundation for state-dependent stimulation protocols that could significantly reduce response variability and enhance therapeutic precision. By addressing the fundamental question of when to stimulate for optimal outcomes, this work provides a pathway toward evidence-based, personalized brain stimulation approaches.

## 2 Results

We analyzed open access data from 36 patients who underwent varying numbers of intracranial electrical stimulation (iES) sessions (ranging from 4 to 17 per patient), each consisting of repeated single-pulse stimulations delivered to clinically defined target regions. Each session included up to 60 stimulation trials, with stimuli separated by 2 seconds and a random jitter of 200–500 ms to reduce anticipatory effects. During each session, brain activity was recorded simultaneously using stereo-EEG (SEEG) and high-density EEG (hdEEG) at the sensor level. Source reconstruction onto 90 cortical regions of interest (ROIs) was performed for supplementary analyses. For each subject, session, and modality, we segmented the data into trials of 1-second duration, aligned to the stimulus onset (-300 ms to +700 ms; *t* = 0; Fig. 1A–B).

**Figure 1:**
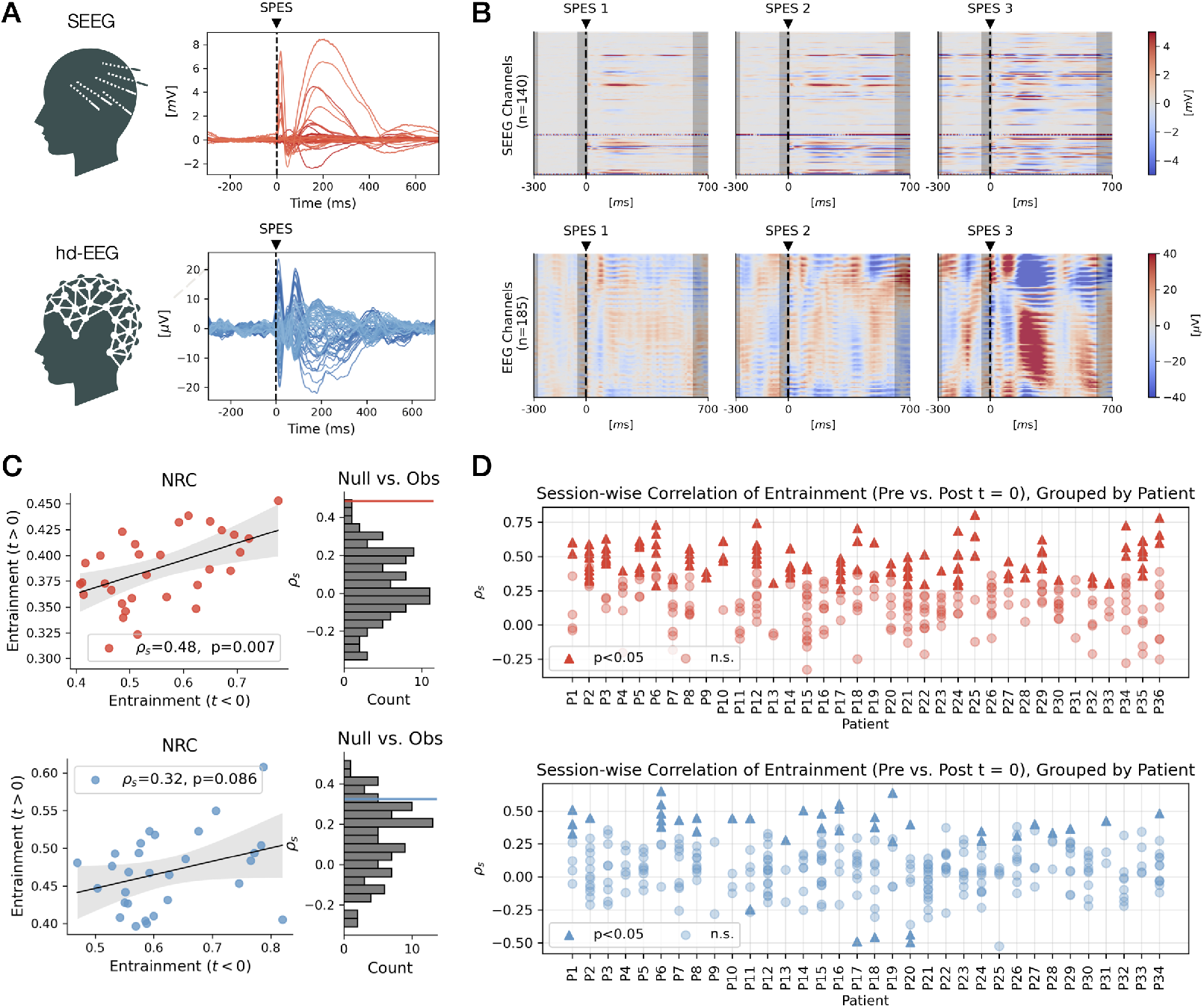
Pre-stimulus functional connectivity predicts post-stimulus network responses. **(A)** Trial-averaged responses to single-pulse electrical stimulation (SPES) in Participant P03, Session S08, recorded via SEEG (top, red) and hdEEG (bottom, blue). Traces are aligned to stimulus onset (black dashed line) and represent mean activity across all channels. **(B)** Voltage dynamics from three individual SPES trials (SPES 1–3) in the same session. Top: SEEG channels (*n* = 140); bottom: hdEEG channels (*n* = 185). Color maps show time-resolved voltage activity relative to stimulation onset, revealing substantial trial-to-trial variability in spatial and temporal patterns. Gray-shaded regions were excluded from pre- and post-stimulus metric computation. **(C)** Example Neural Response Curve (NRC) analysis from a representative session. Left: Scatterplot showing the relationship between Entrainment (mean phase coherence) before (*t <* 0) and after (*t >* 0) stimulation across trials. Right: Histogram comparing the observed Spearman’s correlation (*ρ*_*s*_) against a null distribution from surrogate data. A strong positive *ρ*_*s*_ suggests that pre-stimulus connectivity constrains post-stimulus synchronization. **(D)** Group-level analysis of session-wise Entrainment correlations (*t <* 0 vs. *t >* 0), grouped by patient. Triangles indicate significant correlations (*p <* 0.05), circles indicate non-significance. Top: SEEG (red); Bottom: hdEEG (blue). Across modalities, sessions with higher pre-stimulus Entrainment tended to produce stronger post-stimulus Entrainment, supporting the hypothesis that the ongoing large-scale brain state shapes both the variability and structure of evoked responses.

A common approach to studying the effects of brain responses to stimulation is to rely on averaged event-related potentials (ERPs, sometimes referred to as the ‘deterministic’ response; Fig. 1A). While averaging stimulus responses across many trials increases the signal-to-noise ratio, it also obscures unique information contained in single trials. Here, we analyze single-trial responses under the hypothesis that spontaneous neural dynamics provide trial-specific predictors of evoked responses (Fig. 1B).

### 2.1 Neural Response Curves Reveal Predictive Power of Spontaneous States

To systematically characterize the influence of pre-stimulus neural activities on evoked responses, we defined a set of Metrics of Interest (MOIs), each capturing distinct aspects of large-scale neural dynamics. For each trial, MOIs were computed in two non-overlapping windows: ([*−*300, *−*50] ms) pre-stimulus and ([30, 700] ms) post-stimulus. This approach avoided contamination from stimulus-related artifacts and autocorrelated noise near *t* = 0.

For each MOI pair, we constructed a Neural Response Curve (NRC) by plotting the pre-stimulus metric against the corresponding post-stimulus metric across trials. Representative NRCs from one session across modalities (Fig. 1C, left) show strong positive correlations between pre- and post-stimulus *entrainment*, as measured by mean phase coherence across brain regions (SEEG: Spearman’s *ρ*_*s*_ = 0.48, *p* = 0.006; hd-EEG: *ρ*_*s*_ = 0.32, *p* = 0.08). In other words, trials starting with low (high) pre-stimulus entrainment tended to result in low (high) evoked entrainment.

To test whether these correlations reflected genuine neural dynamics rather than random fluctuations, we generated surrogate datasets by shuffling channel assignments across trials (i.e., reassigning each signal from one trial to another randomly selected trial). This preserved both trial-averaged data and within-channel autocorrelation, but disrupted inter-channel dependencies. Thus, observed and surrogate datasets were equivalent in their deterministic (trial-averaged) event-related potential responses but differed in their trial-specific network dynamics and NRCs.

For each session, statistical significance was assessed by comparing the observed pre–post correlation to the distribution of correlations from surrogate datasets (e.g., Fig. 1C, right), with the *p*-value defined as the proportion of surrogate correlations more extreme than the observed value.

At the single-patient level, several NRCs from the real data showed significantly stronger pre–post correlations than surrogates (Fig. 1D; triangles indicate sessions with *p <* 0.05 and circles indicate non-significant sessions), indicating that spontaneous large-scale dynamics exert a measurable influence on evoked response variability.

### 2.2 Identifying Robust Predictive Features Among Metrics of Interest

To identify informative features among the 125 MOIs tested, we selected five representative metrics: Salience, Peak Synchrony, Entrainment, Fluidity, and Complexity (Fig. 2 A; see Methods). These metrics were selected based on their interpretability and consistent performance across sessions, and span a range of theoretical constructs, including standard signal processing, synchrony, network dynamics, and signal diversity measures.

**Figure 2:**
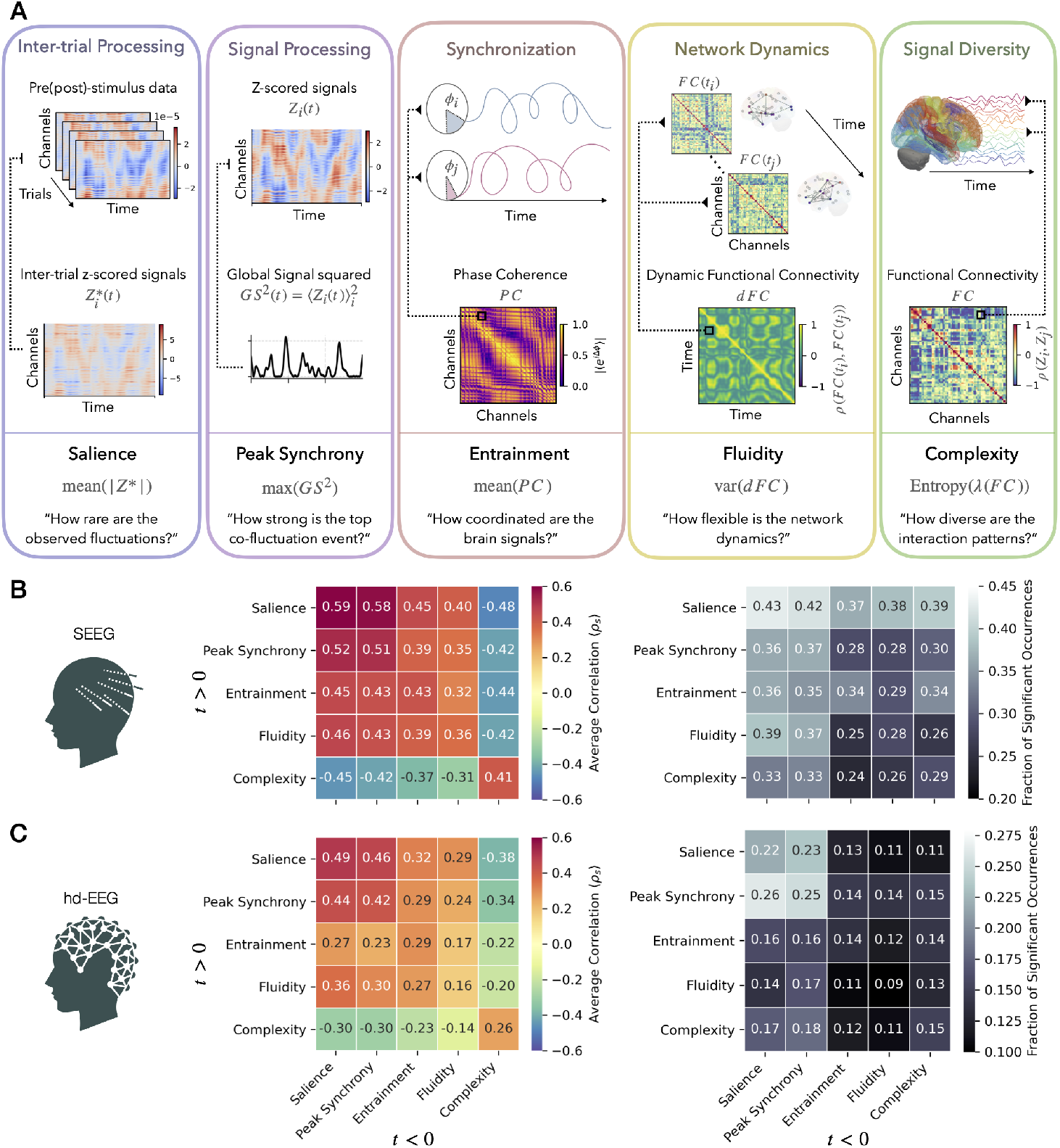
Pre-stimulus brain states predict post-stimulus responses across multiple functional metrics: **(A)** Schematic representation of the analytical pipelines for the five metrics of interest (MOIs) selected among the 125 MOIs analyzed. Each colored box illustrates how the metric is extracted (see Methods section for details), along with its theoretical interpretation. **(B)** SEEG data. Left: average Spearman correlation (*ρ*_*S*_) between metric values computed before stimulation (*t <* 0; x-axis) and after stimulation (*t >* 0; y-axis), across all pairwise combinations of the five metrics. Right: fraction of sessions in which each metric pair showed a significant correlation (*p <* 0.05) compared to null distributions. **(C)** hd-EEG data, same format as in (B). While overall effect sizes and significance rates are reduced compared to SEEG, the qualitative pattern of pre → post metric correlations is preserved. Similar results for source-reconstructed hd-EEG data are shown in Supplementary Figure S5 A. Overall, a diverse feature array demonstrates that spontaneous brain activity modulates how the brain will respond to stimulation, and these predictive relationships generalize across recording modalities.

We examined the relationship between pre- and post-stimulus values of these MOIs using both SEEG and hd-EEG data. For each session and MOI pair, we computed the Spearman correlation coefficient between pre- and post-stimulus values. The left-hand panels in Fig. 2B-C show, for each metric pair, the average correlation across sessions in both modalities. Next, we compared the observed correlation to a null distribution generated from 100 surrogate datasets (as in Fig. 1D). The right-hand panels in Fig. 2B-C show, for each metric pair, the fraction of sessions in which the observed correlation exceeded the null expectation at a two-tailed threshold of *p <* 0.05 (uncorrected).

These results show that several features of spontaneous brain activity predict the structure of post-stimulus responses. For example, a consistent pattern is that evoked responses become less complex when stimulation occurs during periods of high synchrony in ongoing activity. To assess the generality of these effects, we repeated the same analysis for all 125 MOIs (see Supplementary Figs. 1–4). While several metrics showed robust and consistent predictive value across sessions, others exhibited limited or inconsistent performance, underscoring that not all MOIs are equally informative for predicting evoked responses.

Although the absolute strength of correlations was higher in SEEG (likely due to its superior spatial resolution), the overall pattern of significant MOI pairs was preserved in the hd-EEG data (Spearman’s correlation between the left matrices in Fig. 2B-C: *ρ*_*s*_ = 0.93, *p <* 0.001). This cross-modal consistency supports the robustness and generalizability of our findings.

### 2.3 Stimulation reduces variability across trials

We first quantified how stimulation affected the variability of neural responses across trials. For each participant and session, we measured the trial-to-trial standard deviation (STD) of both pre- and post-stimulation values across the five representative metrics (Salience, Peak Synchrony, Entrainment, Fluidity, and Complexity; Fig. 3A–B).

**Figure 3:**
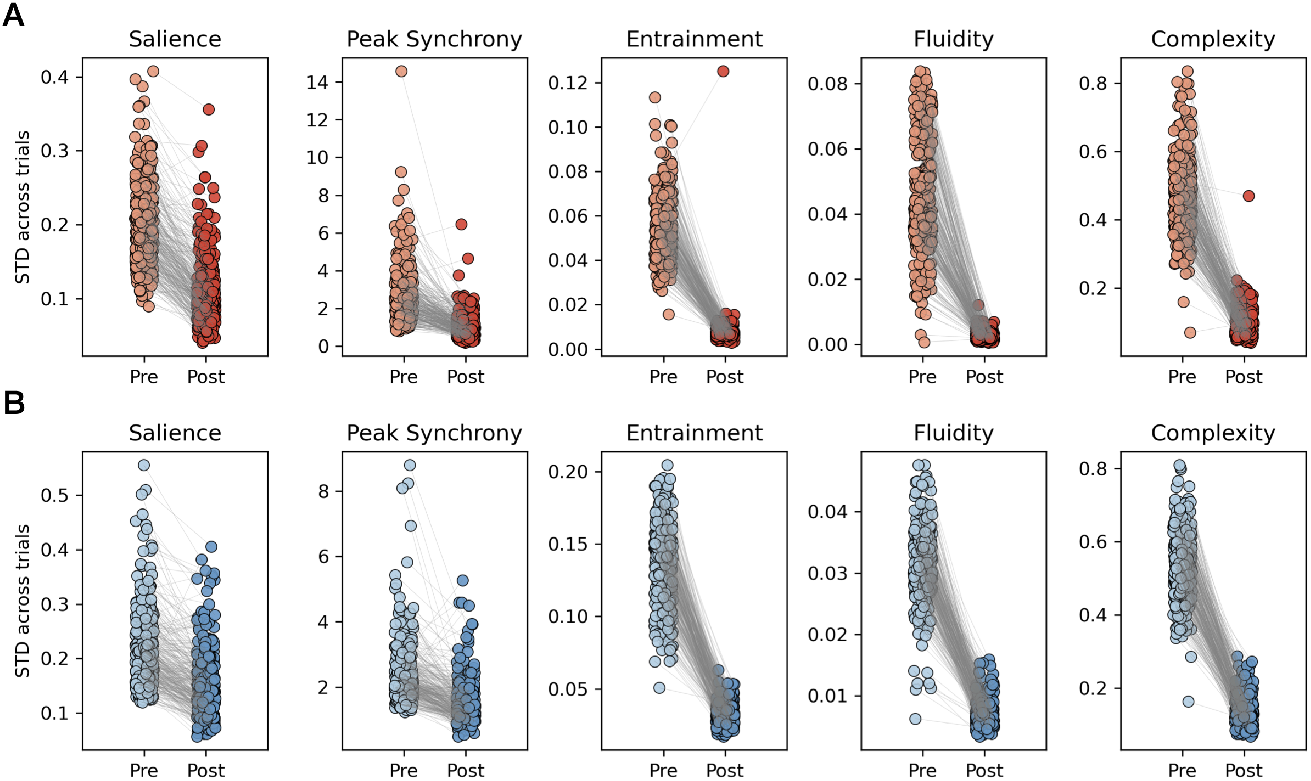
Stimulation reduces trial-to-trial variability across metrics and modalities. (A) Stereotactic EEG (SEEG). (B) High-density EEG (hd-EEG). For each participant and session, the standard deviation (STD) of pre- and post-stimulation values was computed across trials for five representative metrics: Salience, Peak Synchrony, Entrainment, Fluidity, and Complexity. Each dot represents one participant–session; gray lines connect pre- and post-stimulation values from the same participant and session. Across all metrics and modalities, post-stimulation responses consistently exhibit reduced variability compared to pre-stimulation activity. These results show that while stimulation robustly constrains neural responses, substantial variability persists.

Across modalities and metrics, stimulation induced a robust and highly significant reduction in variability. In SEEG (Fig. 3A), mean STD values decreased markedly after stimulation: for Complexity from 0.47 to 0.10, for Peak Synchrony from 2.65 to 1.01, for Salience from 0.20 to 0.11, for Fluidity from 0.046 to 0.003, and for Entrainment from 0.054 to 0.008 (all *p <* 10^*−*50^, Wilcoxon signed-rank test). Comparable effects were observed in hd-EEG (Fig. 3B), where STD reductions were found for Complexity (0.54 to 0.14), Peak Synchrony (2.40 to 1.47), Salience (0.22 to 0.15), Fluidity (0.031 to 0.007), and Entrainment (0.14 to 0.033; all *p <* 10^*−*45^). Together, these results establish that stimulation consistently reduces variability of neural responses across participants, sessions, metrics, and modalities. This provides direct evidence that stimulation drives brain dynamics into more constrained regimes.

Importantly, however, the overall variability remained substantial even after stimulation. Pre-stimulus values often reached very high levels (e.g., mean STD = 2.6 for Peak Synchrony in SEEG; in hd-EEG), and considerable dispersion persisted in post-stimulus responses. This highlights that spontaneous variability is not merely noise but reflects meaningful fluctuations in brain state. As such, uncontrolled stimulation necessarily interacts with a broad distribution of ongoing neural dynamics, contributing to heterogeneous outcomes. These findings highlight the importance of explicitly incorporating pre-stimulus activity into stimulation strategies to further reduce variability.

### 2.4 Conditioning on pre-stimulus states further reduces variability

We next asked whether the stabilization induced by stimulation could be further enhanced by taking into account the ongoing pre-stimulus state. To this end, we compared the variability of post-stimulation responses under two conditions: (i) when only trials in the lowest quartile of pre-stimulus values (Q1) were included, and (ii) when an equally sized random subset of trials was selected.

Figures 4A–B illustrate this procedure for the Peak Synchrony → Peak Synchrony pair in one representative SEEG (panel A) and hd-EEG (panel B) session. Constraining trials to the Q1 subset yielded visibly tighter post-stimulus responses (left), compared to broader dispersions under random selection (right). The standard deviation of post-stimulus values (colored label) was consistently lower for Q1 trials. Figures 4C–D show the global signals (channel-averaged activity) for each of the Q1 and the random trials. Figures 4E–F display the standard deviation of the global signals across Q1 (continuous line) and random trials (dashed line), with the latter demonstrating higher inter-trial variability.

**Figure 4:**
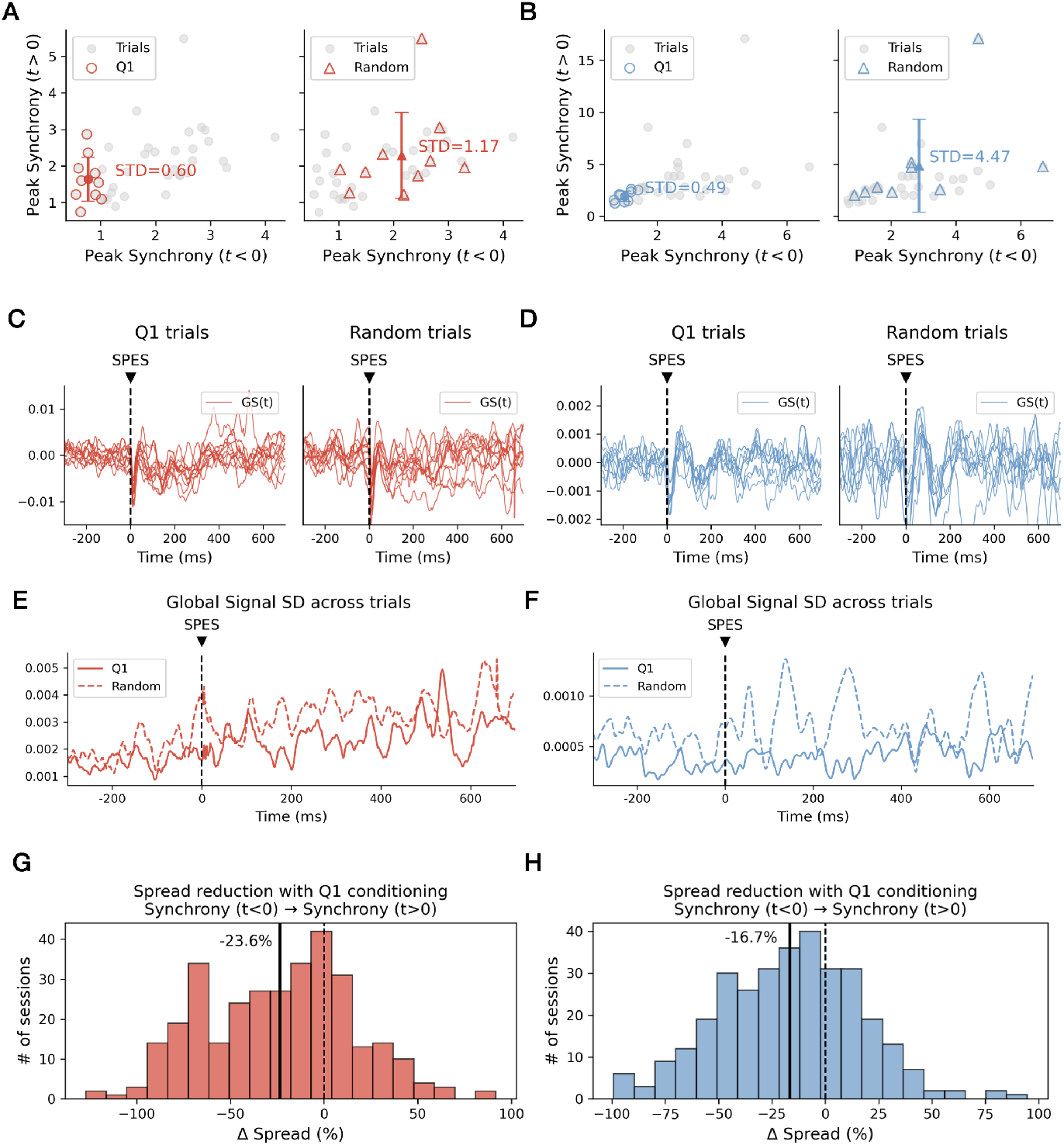
Conditioning on pre-stimulus states reduces variability of stimulation responses. (A–B) Example session (P1, S1) for the Peak Synchrony → Peak Synchrony pair in SEEG (A) and hd-EEG (B). Left: post-stimulation values when trials were constrained to the lowest quartile of pre-stimulus values (Q1). Right: equally sized random trial subset. Colored labels indicate the post-stimulus spread (STD). (C–D) Global signals (channel-averaged activities) for Q1 trials (left) and randomly selected trials (right) in SEEG (C) and hd-EEG (D). (E-F) Standard deviation of the global signals across Q1 trials (continuous line) and randomly selected trials (dashed line) in SEEG (E) and hd-EEG (F). (G-H) Histograms of Δ Spread values (Q1 – Random, relative to full-trial spread) across all sessions for the same metric pair. Each bar corresponds to one participant–session. Negative values indicate reduced variability under Q1 conditioning; vertical line marks the across-session mean.

To quantify the increased dispersion in random trials compared to Q1 trials across all sessions, we computed the relative spread of post-stimulus values with respect to the full-trial spread, and expressed the difference between Q1 and Random as Δ Spread (see Methods). Panels G–H show the distribution of Δ Spread values across sessions for the same metric pair. The histograms demonstrate a systematic shift toward negative values, indicating that Q1 conditioning reduced variability in the majority of sessions. The vertical solid line denotes the mean reduction across sessions (SEEG: –23.6%, *p* = 8.6 *×* 10^*−*20^; hd-EEG: –16.7%, *p* = 4.3 *×* 10^*−*16^; Wilcoxon signed-rank tests).

The strongest effects were observed when both pre- and post-metrics corresponded to Peak Synchrony, but significant reductions were evident across multiple metric combinations (Suppl. Fig. S7).

Together, these results establish that trial-to-trial variability is not random, but can be systematically reduced by conditioning stimulation on specific pre-stimulus states. Clinically, this suggests that real-time monitoring of ongoing brain dynamics could provide a principled route to improve the reliability and reproducibility of brain stimulation interventions.

### 2.5 Pre-Stimulus Context: Local Versus Global Predictive Power

We next tested whether the spatial extent of pre-stimulus activity influenced the ability to predict evoked responses. To this end, we varied the radius (*R*) of included brain regions, ranging from 5% (local neighborhood around the stimulation site) to 100% (whole-brain activity).

As an illustrative example, Figs. 5A–B show the *Entrainment → Entrainment* NRC correlation for each participant as a function of radius. In SEEG, the correlation increased on average from 0.131 at *R* = 5% to higher values with broader radii, with a significant positive fixed effect of radius in a linear mixed-effects model (*β*_radius_ = 0.002, *p <* 10^*−*85^). In hd-EEG, the same effect was observed (*β*_radius_ = 0.001, *p <* 10^*−*21^). These results indicate that incorporating a greater proportion of the network into the pre-stimulus analysis significantly improves the predictability of post-stimulus functional connectivity patterns.

**Figure 5:**
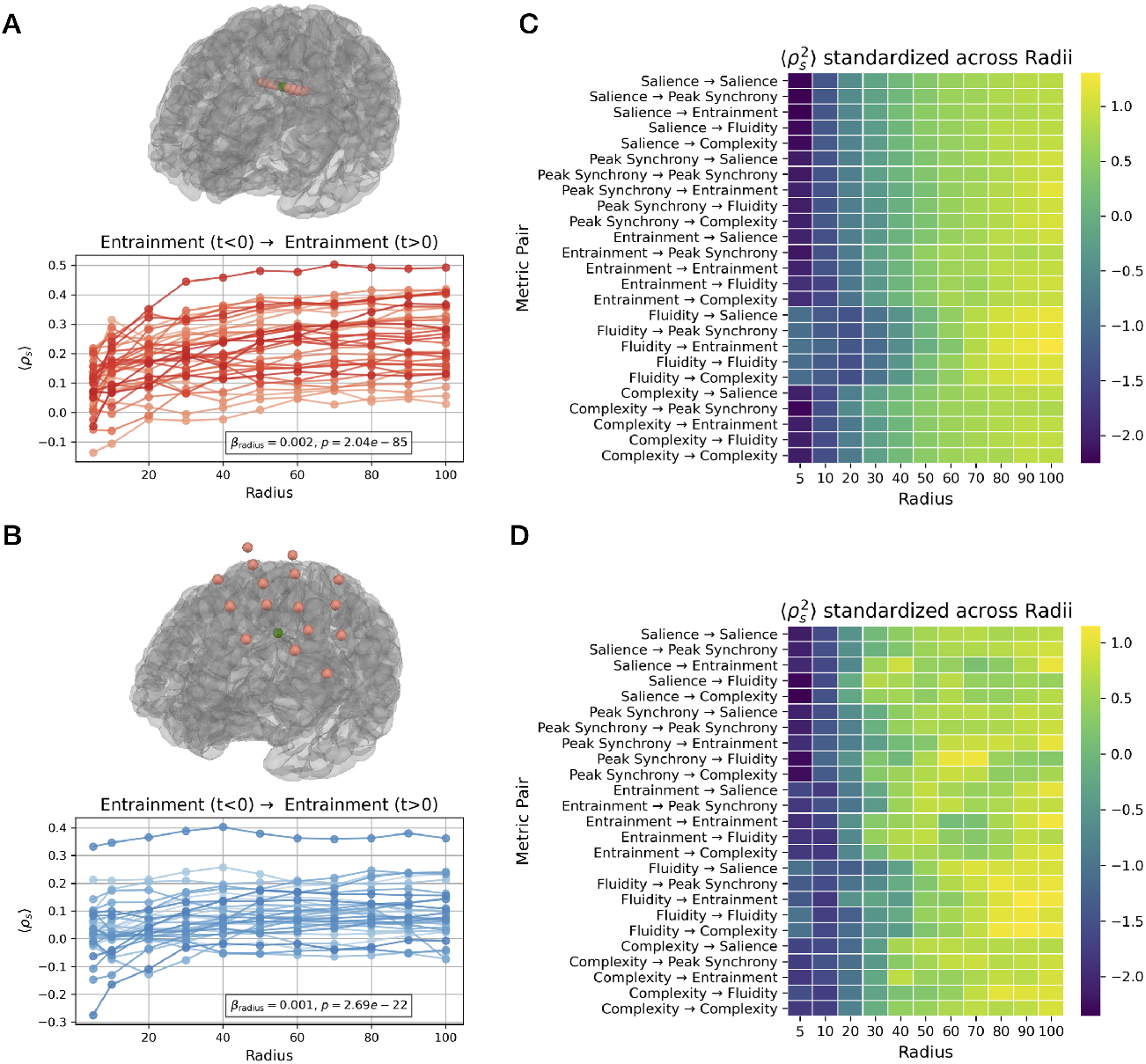
Broader pre-stimulus network dynamics enhance prediction of post-stimulus responses: **(A)** SEEG data: session-wise correlation between pre- and post-stimulus Entrainment as a function of the radius used to compute pre-stimulus connectivity (percentage of channels surrounding the stimulation site). Each line represents one subject. Smaller radii reflect local prestimulus activity, while larger radii incorporate broader network context. **(B)** hd-EEG data: same format as (A). For both modalities, incorporating a greater proportion of the network into the prestimulus analysis significantly improves the predictability of post-stimulus functional connectivity patterns. Linear mixed-effects modeling confirmed a significant positive effect of radius on pre/post Entrainment correlation in SEEG (*β*_radius_ = 0.002, *p* = 2.04 *×* 10^*−*85^) and hd-EEG (*β*_radius_ = 0.001, *p* = 2.69 *×* 10^*−*22^). **(C)** SEEG data: each row represents one pre → post metric pair, each column a radius value. Color denotes the squared correlation 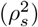 between pre- and post-stimulus values, averaged across sessions and standardized within each metric pair across radii. This highlights the relative gain in ‘predictability’ as a function of radius, regardless of baseline correlation levels. **(D)** hd-EEG data: same format as (C). Across both modalities, larger radii consistently yield higher standardized correlations, indicating that broader pre-stimulus network context enhances the ability to predict post-stimulus connectivity patterns. Similar results for source-reconstructed hd-EEG data are shown in Supplementary Figure S5 B. These findings generalize across multiple metric pairs (see Supplementary Figure S6 for other examples) and confirm that distributed network dynamics before stimulation better explain evoked responses than localized activity alone.

We then extended this analysis to all metric pairs (Figs. 5C–D), where the color maps show standardized correlation values 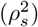 across radii for visualization purposes. Using the non-standardized values, mean correlation across all metric pairs in SEEG increased from 0.0565 at *R* = 5% to 0.1399 at *R* = 100% (+147.6% change; Wilcoxon signed-rank test: *W* = 325.0, *p* = 2.98 *×* 10^*−*8^). In hd-EEG, mean correlation increased from 0.0469 to 0.0631 (+34.6% change; Wilcoxon signed-rank test: *W* = 325.0, *p* = 2.98 *×* 10^*−*8^).

Together, these results demonstrate that distributed pre-stimulus network dynamics—capturing a larger fraction of the brain—are more predictive of evoked responses than local measurements, and that this effect is robust across metrics and recording modalities.

### 2.6 Target Functional Network Shapes Stimulus Effects Predictability

Finally, we investigated whether the functional network identity of the stimulation site modulates the relationship between pre- and post-stimulus activity. Target regions were grouped according to canonical functional networks (as in [30]): default mode (DMN), limbic (LIM), control (CON), dorsal attention (DAN), ventral attention (VAN), sensorimotor (SMN), and visual (VIS).

In SEEG data (Fig. 6A), predictability was highest when stimulation occurred in SMN and VIS regions, while DMN and LIM regions showed substantially lower explained variance. Linear mixed-effects modeling, with patient as a random intercept, confirmed a significant main effect of target network. Using DMN as the reference category, explained variance was significantly higher for CON (*β* = 0.028, *p <* 0.001), VAN (*β* = 0.014, *p* = 0.003), SMN (*β* = 0.028, *p <* 0.001), and VIS (*β* = 0.039, *p <* 0.001) targets, and for LIM (*β* = 0.013, *p* = 0.015), while DAN did not differ significantly from DMN (*β* = 0.012, *p* = 0.061). This pattern suggests that evoked responses are more strongly constrained by pre-stimulus states in regions engaged in direct sensory and motor processing, potentially reflecting more stable or stereotyped functional architectures that render their responses more susceptible to ongoing network dynamics.

**Figure 6:**
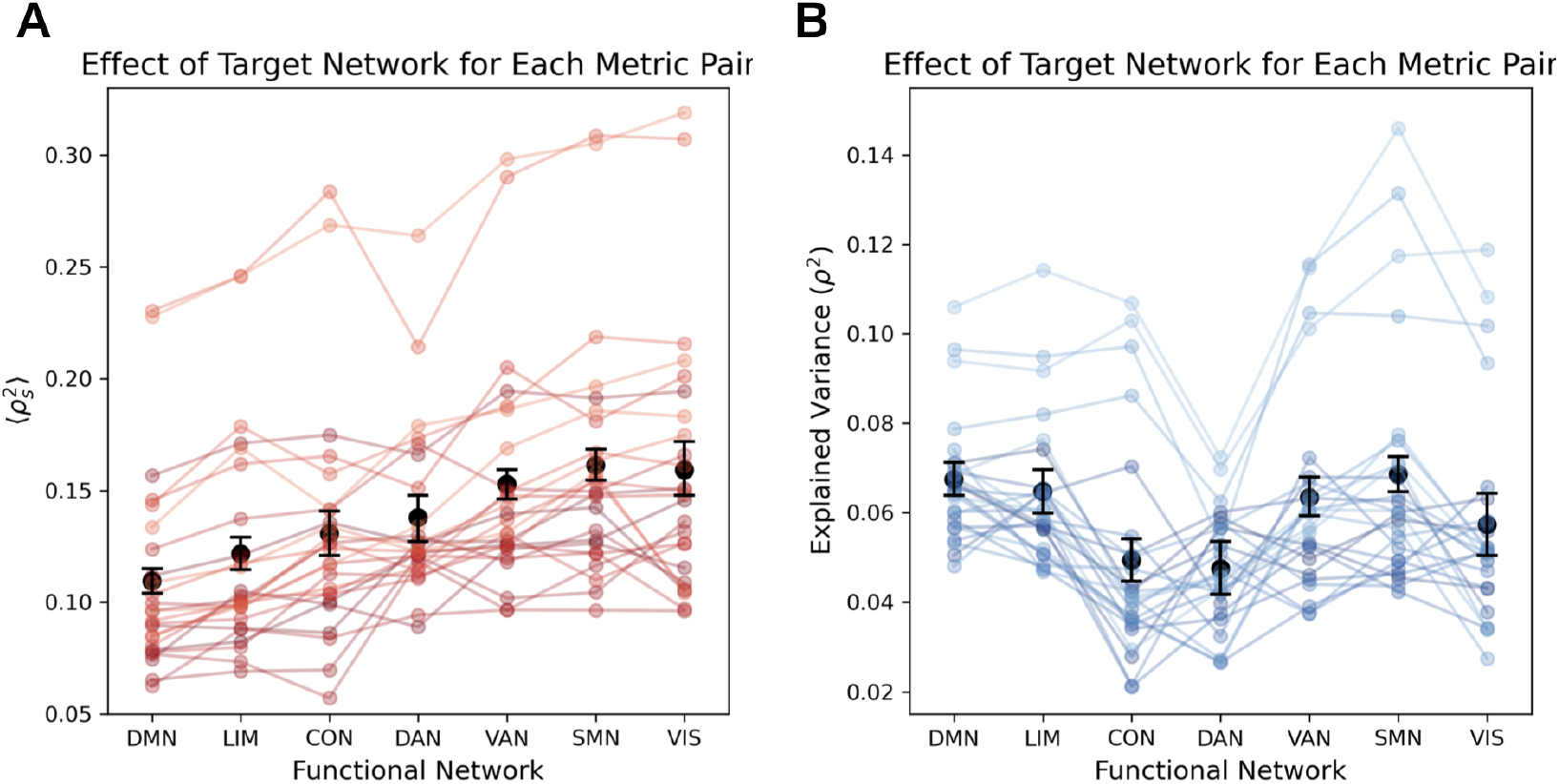
Pre → post predictability varies with stimulation site across functional networks. **(A)** SEEG data: session-averaged explained variance 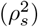 of post-stimulus functional connectivity predicted from pre-stimulus connectivity, grouped by functional network of the stimulation site. Each line represents one metric pair; black error bars indicate mean *±* SEM across all metric pairs. Predictability is highest for sensorimotor (SMN) and visual (VIS) targets, and lowest for default mode (DMN) and limbic (LIM) targets, with LMEM confirming significant network effects. **(B)** hd-EEG data: same format. While the overall magnitude of explained variance is lower and network contrasts differ in direction, LMEM still indicates significant effects of network identity. Similar results for source-reconstructed hd-EEG data are shown in Supplementary Figure S5 C. Overall, the functional network context of the stimulation site influences how strongly spontaneous activity constrains evoked responses, with sensory/motor regions in SEEG showing greater predictability than higher-order association networks, whereas hd-EEG reveals a different network hierarchy.

Hd-EEG data revealed a qualitatively different pattern (Fig. 6B), with overall lower explained variance and opposite directionality for network contrasts. The LMEM indicated significantly lower explained variance compared to DMN for LIM (*β* = *−*0.009, *p* = 0.010), CON (*β* = *−*0.017, *p <* 0.001), DAN (*β* = *−*0.026, *p <* 0.001), VAN (*β* = *−*0.014, *p <* 0.001), SMN (*β* = *−*0.013, *p <* 0.001), and VIS (*β* = *−*0.022, *p <* 0.001). Despite these modality-specific differences, both analyses indicate that network-specific properties influence the degree to which spontaneous activity governs stimulation outcomes.

## 3 Discussion

This study establishes that spontaneous neural activity systematically predicts brain stimulation responses across trials, sessions, and individuals. Using simultaneous stereotactic and scalp EEG recordings from 36 patients and *∼*320 stimulation sessions, we demonstrate that pre-stimulus brain states account for a substantial portion of post-stimulus response variability—explaining on average 24.2% in SEEG and 7.7% in hd-EEG, with predictive power reaching up to 84.2% and 64.3% respectively in individual sessions. These findings fundamentally challenge the treatment of stimulation variability as random noise and reveal structured patterns that could inform future neuromodulation strategies.

Our results demonstrate that spontaneous large-scale neural dynamics serve as robust predictors of stimulus effects, with specific metrics—particularly those reflecting network synchronization, functional connectivity, and signal diversity—consistently showing predictive value across sessions (Fig. 1-2). This extends beyond well-established phase-dependency effects [16] to encompass multiple dimensions of network organization. Phase-dependent local brain states determine the impact of brain stimulation on motor network synchronization [15], and our results substantially expand this framework by demonstrating that complex, multidimensional features of spontaneous activity provide systematic predictive information about stimulation outcomes across multiple brain networks and recording modalities. Moreover, phase-based measures are challenging to extract reliably outside the motor cortex, making our multidimensional approach particularly valuable for broader clinical applications across diverse brain regions.

In addition to this, we found that stimulation consistently reduced trial-to-trial variability across all metrics and recording modalities (Fig. 3), demonstrating that stimulation might drives brain dynamics into more constrained regimes. This supports dynamical systems perspectives viewing ongoing activity as metastable attractor states [26]. However, substantial variability persisted even after stimulation, highlighting that spontaneous fluctuations reflect meaningful brain states rather than mere noise. Critically, conditioning stimulation on specific pre-stimulus states (lowest quartile) reduced post-stimulus variability by an average of 23.6% across sessions compared to random selection (Fig. 4), providing direct evidence that this variability contains exploitable structure that can improve stimulation consistency. This finding further shows that brain stimulation outcomes can be systematically optimized by monitoring and targeting specific spontaneous brain states.

Moreover, we found that the spatial extent of pre-stimulus analysis proved crucial for prediction accuracy. Mean correlation across metric pairs in SEEG increased from 0.0565 at R = 5% to 0.1399 at R = 100% (+147.6% change), while in hd-EEG correlations increased from 0.0469 to 0.0631 (+34.6% change) when considering whole-brain versus local pre-stimulus activity (Fig. 5). This dramatic improvement demonstrates that distributed pre-stimulus network dynamics are more predictive of evoked responses than local measurements alone, supporting the view of the brain as a complex dynamical system where global states modulate local responses. This distributed perspective aligns with computational models showing that stimulation responses reflect network-wide reverberation patterns rather than purely local effects [33], and with evidence that network-level macroscale structural connectivity predicts the propagation pattern induced via brain stimulation [34, 35].

Finally, our findings showed that the functional network identity of stimulation targets significantly influenced predictability patterns (Fig. 6). Specifically, sensorimotor regions showed the highest predictability in SEEG, yielding approximately 1.2-fold higher explained variance than association networks. The dependence of the response with respect to a core-periphery axis has been shown both theoretically [36], and validated empirically targeting regions V1 (bottom of the visual hierarchy) and the frontal eye fields (top of the visual hierarchy) [37]. These variations might reflect fundamental differences in functional architecture across the cortical hierarchy [38]. Sensorimotor regions, anchored in stereotyped input-output transformations [39], exhibit more deterministic state-response relationships compared to higher-order association networks that integrate diverse and temporally extended inputs [40]. This pattern also aligns with recent evidence that stimulation mapping reveals gradients of excitability and recurrence in cortical networks, with high-order networks showing greater dependence on recurrent feedback [30].

Overall, these findings have direct implications for improving clinical brain stimulation protocols. Variability in stimulation outcomes currently limits therapeutic efficacy across neurological and psychiatric disorders [12, 41–44]. Our identification of robust predictive metrics—including Salience, Peak Synchrony, Entrainment, Fluidity, and Complexity—provides concrete targets for real-time state estimation in closed-loop stimulation systems [45–47]. Current implementations often rely on simple local markers such as the alpha phase in target regions [48], but our systematic evaluation of 125 metrics reveals that multidimensional features of spontaneous activity offer superior predictive power. These metrics likely reflect neurophysiological states associated with integration, excitability, and sensitivity to perturbations, making them strong candidates for clinical implementation.

Translation to clinical practice appears feasible given current technology. Whole-brain analysis requires processing latencies under 100ms, which are achievable with contemporary hardware. Cross-modal validation between invasive and non-invasive recordings suggests that EEG-based implementations can capture sufficient predictive information for real-time applications. In disorders of consciousness, where stimulation complexity serves as a consciousness biomarker [49–51], timing stimulation to optimal pre-stimulus states could evoke more reliable responses with fewer trials, improving diagnostic efficiency. Similarly, for treatment-resistant depression, epilepsy, and other conditions, state-dependent stimulation could enhance therapeutic outcomes by reducing current trial-and-error approaches [52, 53]. In Parkinson’s disease, it may further suggest strategies to guide pharmacological therapy, aiming to place the brain in a dynamical regime where electrical stimulation is maximally effective [54, 55]. Addressing these aspects could help overcome current limitations, and help explain variable clinical responses and recent negative trials [56]. From a theoretical standpoint, computational models showed that perturbation effects depend more on dynamical brain states than static connectivity patterns [57, 58].

These findings extend beyond electrical stimulation, resonating with state-dependent perception research [59–63]. This convergence supports a unifying principle: the brain’s momentary state governs responsiveness to both therapeutic interventions and natural sensory inputs, emphasizing the fundamental importance of timing in neuromodulation strategies.

Several limitations should be considered when interpreting these findings. Our patient population consisted of individuals with epilepsy, and generalizability to other clinical populations requires validation. While we tested 125 metrics across multiple domains, optimal feature combinations for real-time implementation remain to be determined. Our analysis focused on single-pulse stimulation; extending to complex stimulation protocols presents additional challenges. Finally, computational requirements for real-time whole-brain analysis may limit clinical feasibility in some settings.

Overall, these results provide a quantitative foundation for moving beyond empirical approaches toward predictive frameworks for brain stimulation. By revealing that stimulation variability contains structured, exploitable patterns rather than representing random noise, this work establishes state-dependent stimulation as a fundamental principle that could significantly improve the reliability and effectiveness of neuromodulation approaches through real-time monitoring and conditioning on optimal brain states.

## 4 Materials and Methods

### 4.1 Participants

The data used in this study were taken from an open dataset collected at the Claudio Munari Epilepsy Surgery Center, Milan [64], where SEEG and scalp hd-EEG were recorded following single-pulse intracerebral electrical stimulation (iES) on 36 patients (median age: 33 *±*8 years; 21 female). All subjects had a history of drug-resistant, focal epilepsy, and were candidates for surgical removal/ablation of the seizure onset zone (SOZ). For details regarding the data acquisition and the preprocessing steps, please refer to the original papers [31, 32].

### 4.2 Data preprocessing

#### Single Pulse Electrical Stimulation

During simultaneous high-density EEG (hd-EEG) and stereo-EEG (SEEG) recordings, single biphasic electrical pulses (positive-negative) were delivered between pairs of adjacent stereotactic contacts on the same electrode, as described in [30–32]. These stimulations were administered with an inter-stimulus interval of 2 seconds and a random jitter of 200–500 ms to reduce anticipatory effects. Brain activity was continuously recorded from all remaining SEEG contacts and from the 256-channel scalp hd-EEG system. Stimulation sessions were included in the analysis according to the following criteria: (i) stimulation was delivered through a bipolar contact located outside the seizure onset zone (SOZ), identified by pathological electrical activity and confirmed retrospectively through post-surgical evaluation; (ii) the stimulated contact did not exhibit spontaneous interictal epileptiform activity, as verified by visual inspection; and (iii) stimulation did not elicit muscle contractions, somatosensory sensations, or cognitive effects. After preprocessing, the number of sessions per subject ranged from 4 to 17 (mean*±*SD: 9 *±* 4), yielding a total of 323 sessions for SEEG and 318 for hd-EEG. Each stimulation session consisted of 15–60 trials for SEEG and 9–57 consecutive trials for hd-EEG. In total, we analyzed 11,070 single stimulations for SEEG and 10,088 for hd-EEG.

#### Functional brain networks

For each trial, the electrical stimulation was delivered to the same anatomical site. The site was then mapped to one of the canonical large-scale functional brain networks [65], allowing each trial to be considered as a unique pairing between the stimulation site and the corresponding stimulated network. The functional networks considered in this study are the default mode network (DMN), dorsal attention network (DAN), ventral attention network (VAN), sensorimotor network (SMN), visual network (VIS), limbic network (LIM), and control network (CON). Network assignment was based on the anatomical location of the stimulated contacts using a standardized brain atlas and parcellation scheme, as in [30].

#### Computational Analysis of Pre- and Post-Stimulus Neural Activity

To investigate the effects of intracranial stimulation on brain dynamics, we analyzed neural data recorded before and after stimulation using a custom computational pipeline implemented in Python, available at the GitHub repository https://github.com/grabuffo/State_Dependent_Brain_Stimulation.git. For each analysis, the time windows corresponding to the pre-stimulus and post-stimulus periods were manually defined. The windows were used to extract segments of interest for subsequent analysis.

#### Feature Extraction

To characterize both pre- and post-stimulus activity, we computed a range of features spanning multiple perspectives, from network-theoretic measures to signal diversity indices. For post-stimulus signals, all metrics were derived from the complete set of available channels. For pre-stimulus signals, metrics were computed from a subset of channels determined by their spatial proximity to the stimulation site, defined as a percentage *R* of channels (e.g., *R* = 5 indicates the 5% nearest channels). For *R* = 100, pre-stimulus metrics were calculated from the full set of channels. The complete set of explored metrics is reported in the open-access repository https://github.com/grabuffo/State_Dependent_Brain_Stimulation.git. Below, we describe the five representative metrics of interest (MOIs) selected for their interpretability and consistent performance across sessions:

1. **Salience** Neural responses to stimulation depend on the baseline excitability state of target regions. Salience quantifies how much brain regions’ activity deviates from their typical patterns, reflecting heightened local excitability or unusual network configurations. Computationally, we measured *salience* as

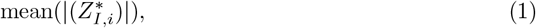

where 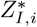 represents the i-th channel signal in the *I*-th trial, standardized relative to all other trials. For each channel and trial, this inter-trial standardization is obtained by standardizing each channel’s activity relative to the mean and standard deviation calculated across all trials of the same window type (pre- or post-stimulus). For example, when analyzing the *I*-th trial in the post-stimulus window (*t >* 0), we subtracted from each channel its mean value calculated across *all* post-stimulus trials and divided by the standard deviation computed from those same windows. Pre- and post-stimulus trials were always processed separately, ensuring that spontaneous and evoked activity were each evaluated against their own variability. High salience indicates the brain is in an atypical state that may respond differently to external perturbation, while low salience suggests a more ‘typical’ baseline from which stimulation effects may be more predictable.
2. **Peak Synchrony** Peak Synchrony identifies moments of maximal global neural coordination, when the largest number of brain regions simultaneously exhibit correlated activity. For each trial and for each window type (pre- or post-stimulus), we standardized each channel’s activity using the normal z-score transformation to obtain the z-scored signals *Z*_*i*_(*t*). We then averaged these z-scored signals across channels to compute the global signal (GS) for that trial and window type. The *peak synchrony* was defined as

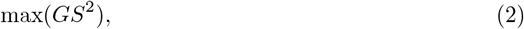

marking the strongest co-fluctuation event observed within the trial. Neurobiologically, these events reflect periods when the brain transitions into highly organized states, similar to the large-scale synchronization observed during cognitive tasks, attention shifts, or pathological events like seizures (though at much smaller magnitudes). Peak synchrony moments may represent critical transition points where the brain is most susceptible to external perturbations. High peak synchrony suggests the brain is operating in a highly coordinated but potentially rigid state, while low peak synchrony indicates more independent regional activity that might be more flexible but less integrated.
3. **Entrainment** Entrainment quantifies network-wide neural synchronization, the degree to which spatially distributed brain regions oscillate in temporal coordination. For each trial and for each window type (pre- or post-stimulus), we applied the Hilbert transform to obtain the instantaneous phase of each channel *ϕ*_*i*_(*t*). We then computed the phase coherence (PC) matrix across all channel pairs

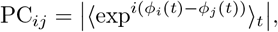 The *entrainment* was defined as the mean value of the upper triangular part of this matrix i.e.,

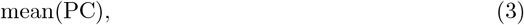

where we exclude the diagonal elements from the mean function. This metric quantifies the degree of network-wide synchrony and reflects the notion that synchronized neuronal populations can mutually entrain each other’s activity. Neurobiologically, synchronized oscillations facilitate information integration across cortical areas and reflect the binding of distributed neural processes into coherent functional networks. Higher entrainment indicates that neural populations are operating in a coordinated state, potentially reflecting enhanced readiness for information processing or reduced capacity for flexible reconfiguration.
4. **Fluidity** Fluidity captures the temporal evolution of functional brain networks, how patterns of inter-regional communication change over time. For each trial and for each window type (pre- or post-stimulus), we first computed the dynamic functional connectivity (dFC) using an approach based on edge co-fluctuation events [66, 67], which allows for time-resolved dFC estimation without sliding windows. For each pair of standardized channel activities *Z*_*i*_(*t*) and *Z*_*j*_(*t*), we defined the edge co-fluctuation signal as

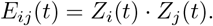 At a fixed time *t*_*a*_, the co-activation pattern defines an instantaneous functional connectivity matrix *FC*_*ij*_(*t*_*a*_) *≡ E*_*ij*_(*t*_*a*_). The dFC between two time points *t*_*a*_ and *t*_*b*_ was then defined as the Pearson correlation between their instantaneous co-activation patterns:

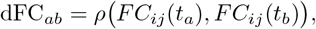

where the correlation is computed over all edges *{ij}. Fluidity* was defined as

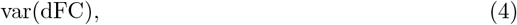

calculated on the upper triangular part of the matrix. This reflects the brain’s capacity for dynamic reconfiguration between different functional states. High fluidity indicates that the brain is transitioning between multiple network configurations, suggesting cognitive flexibility and adaptive capacity. Low fluidity suggests the brain is ‘stuck’ in stable patterns, which may indicate either focused processing or reduced adaptability to perturbation
5. **Complexity** Complexity quantifies the informational richness of brain network organization, how diverse and balanced the patterns of inter-regional communication are. For each trial and for each window type (pre- or post-stimulus), we computed the Pearson correlation matrix between all pairs of channel signals, yielding the Functional Connectivity (FC) matrix for that trial. We then obtained the eigenvalues *{λ*_*k*_*}* of the FC matrix and normalized them such that ∑_*k*_ *λ*_*k*_ = 1. The *complexity* was defined as the Shannon entropy of the normalized eigenvalue distribution [68]

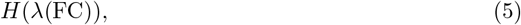

where

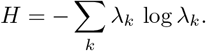 The eigenvalues *λ* of the FC matrix describe the distribution of variance explained by different network modes. Entropy *H* applied to this spectrum quantifies the level of informational richness in the network. This reflects a fundamental principle in neuroscience: healthy brain function requires a balance between integration (coordinated activity) and segregation (specialized processing). High complexity indicates the brain is operating in a rich, multidimensional state with many different network modes active simultaneously, characteristic of conscious, flexible cognition. Low complexity suggests the brain is dominated by a few simple patterns, which may indicate either highly focused processing or pathological states. This metric is particularly relevant for disorders of consciousness, where reduced complexity correlates with diminished awareness

#### Comparison Between Pre- and Post-Stimulus Metrics

To quantify the relationship between pre- and post-stimulus activity, Spearman’s rank correlation *ρ*_*S*_ was computed for each pair of features extracted before and after stimulation. The resulting correlation matrix and corresponding p-value matrix captured the relationships between pre- and post-stimulus neural features.

To evaluate the statistical significance of stimulation-induced effects, we generated a null dataset by independently shuffling the channels across the trials within each session. This procedure breaks the temporal correspondence across channels within single trials, while preserving the temporal structure within each channel and the trial-average response of each channel. This results in a surrogate dataset with identical marginal distributions and signal properties within each channel, but where trial-level synchrony or cross-channel dependencies related to the effects of the stimulation are eliminated. The trial-shuffled null model served as a reference distribution for the observed correlations between MOIs pairs in pre- versus post-stimulus windows. By comparing the values obtained from real data against those from the null distribution, we can identify the effects of stimulation that are above chance level.

#### Quantification of state-dependent variability

For each participant and session, the variability of post-stimulation responses was quantified as the standard deviation (STD) across trials. We compared this variability under two conditions: (i) when trials were restricted to the lowest quartile of pre-stimulus values (Q1), and (ii) when an equally sized subset of trials was randomly selected.

For a given pre–post metric pair, the relative spread was defined as

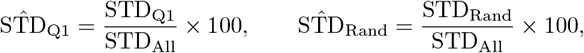

where STD_All_ denotes the variability computed across all trials in that session.

The effect of conditioning was quantified as the difference

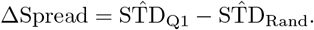

In Fig. 4G–H, negative values of ΔSpread indicate that selecting trials with low pre-stimulus values systematically reduces the variability of post-stimulation responses compared to chance selection. Values were computed for each participant–session and then averaged across sessions to obtain the group-level pre–post metric matrices shown in Fig. S7.

## Acknowledgments

This work was supported by the Marie Sklódowska–Curie Postdoctoral Fellowship (Project CAERUS) under the European Union’s Horizon Europe research and innovation programme, grant agreement No. 101199894. Centre de Calcul Intensif d’Aix-Marseille is acknowledged for granting access to its high-performance computing resources. M.A. was supported by the Excellence Initiative of Aix-Marseille Université – AMidex, a French ‘Investissements d’Avenir’ programme (AMX-21-IET-017).

## Author Contributions

Conceptualization: G. R., P. S., D. M.; Methodology / Analysis: G. R., M. A., D. M.; Writing – Original Draft: G. R., M. A., P. S., D. M.; Writing – Review / Editing: G. R., M. A., G. D., T. F., P. S., D. M.

## Data Availability

As noted above, SEEG and hd-EEG data were obtained from the EBRAINS Knowledge Graph (https://ebrains.eu/) and are also available at the Open Science Framework [64]. The dataset is provided in BIDS format [69] and includes: simultaneous hd-EEG and SEEG from a total of 323 iES sessions, obtained from 36 subjects [31, 32]. In addition, it includes the spatial locations of the stimulating contacts in native MRI space, MNI152 space and Freesurfer’s surface space, as well as the digitized positions of the 185 scalp hd-EEG electrodes. It also contains the MRI of each subject, de-identified with AnonyMi [70].

## Code Availability

Full code for the reproduction of the data analysis described in this paper is freely available online at the repository https://github.com/grabuffo/State_Dependent_Brain_Stimulation.git.

## Supplementary Materials

**Figure S1:**
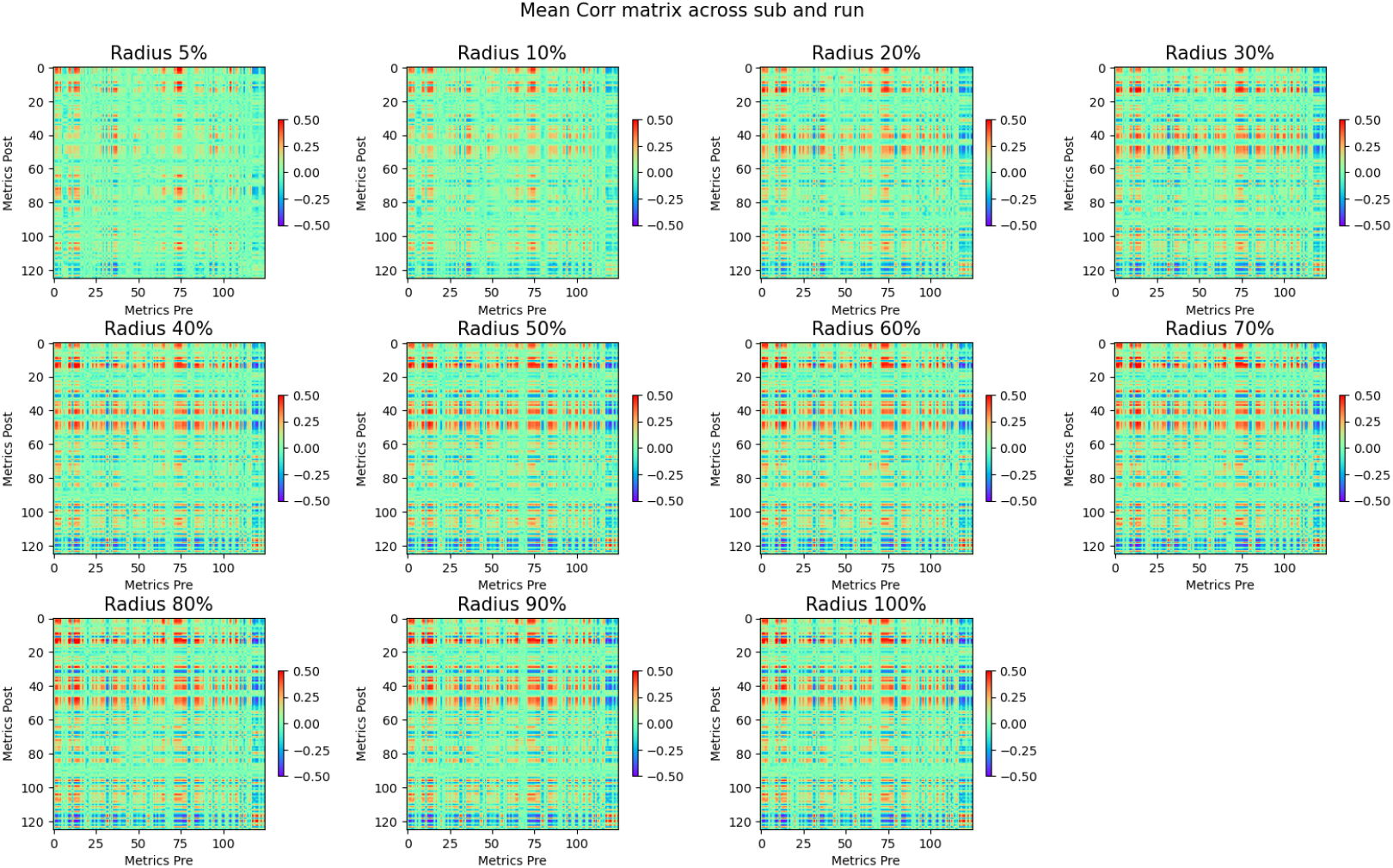
SEEG data: Sessions-averaged correlation matrices for the full set of MOIs (extension of Fig.2), for several values of the radius (defining which channels around the stimulation target were used for measuring the pre-stimulus dynamics). Several (but not all) MOI pairs display a strong correlation between pre- and post-stimulus dynamics. Increasing the radius increases the average correlations across metrics pairs.

**Figure S2:**
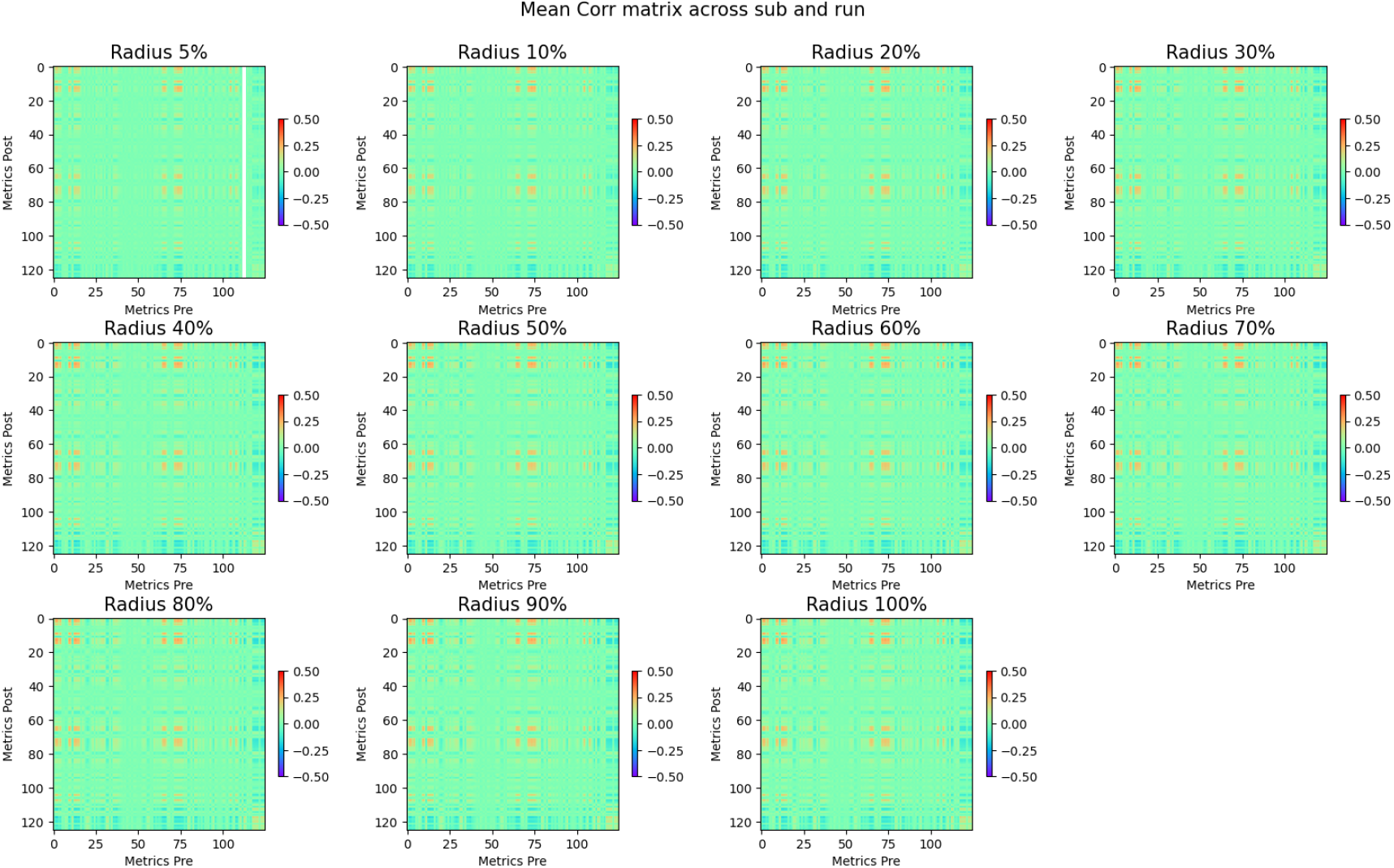
HDEEG sensor level data: Sessions-averaged correlation matrices for the full set of MOIs (extension of Fig.2), for several values of the radius (defining which channels around the stimulation target were used for measuring the pre-stimulus dynamics). Several (but not all) MOI pairs display a strong correlation between pre- and post-stimulus dynamics. Increasing the radius increases the average correlations across metrics pairs.

**Figure S3:**
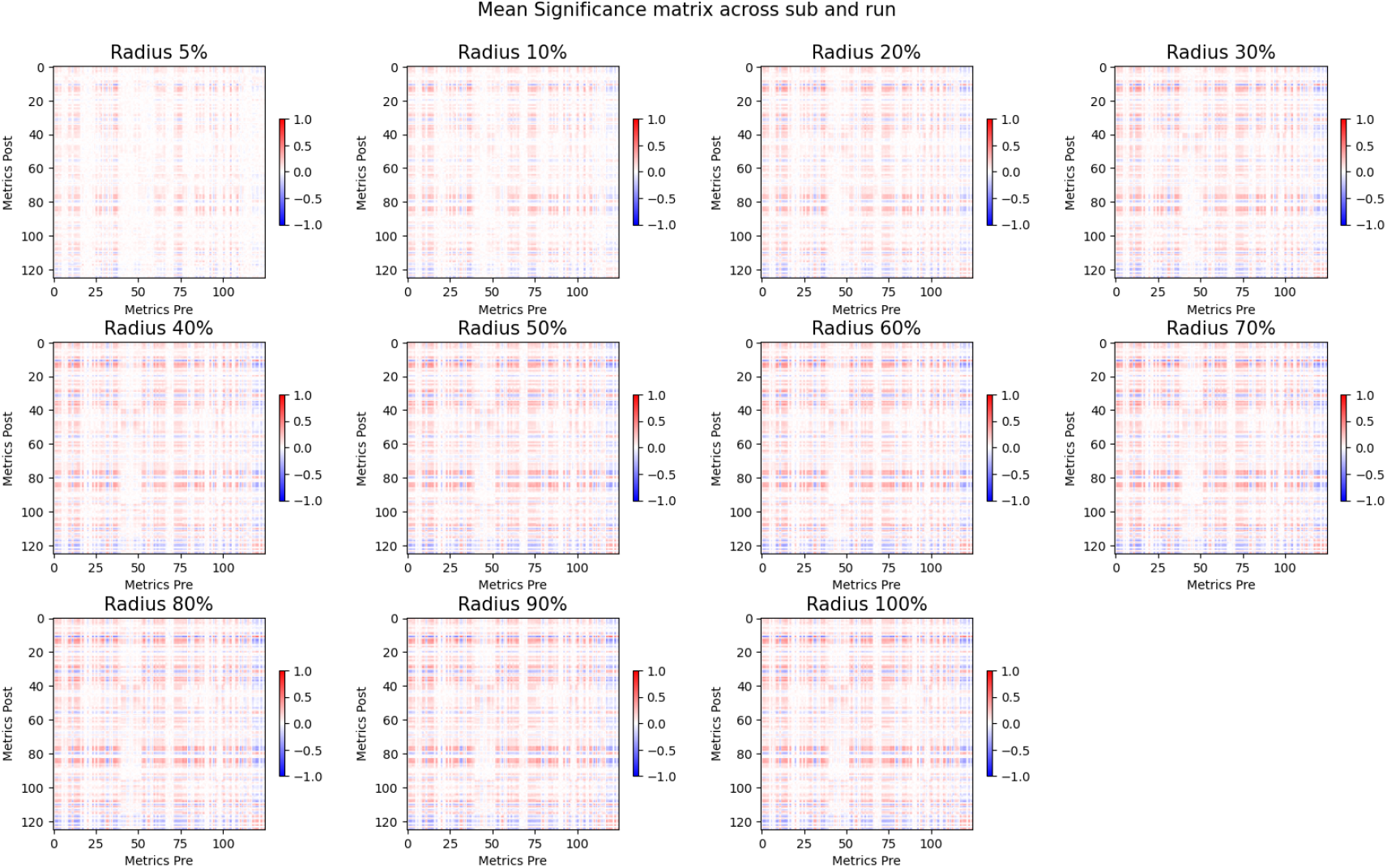
Significance matrices for the full set of MOIs, averaged across all sessions, for varying radii. The matrices’ entries represent the fraction of sessions where the pre/post correlations were significant. Red (blue) represents values larger (smaller) than expected compared to the null model. Several (but not all) MOI pairs display significant levels of correlation between pre- and post-stimulus dynamics.

**Figure S4:**
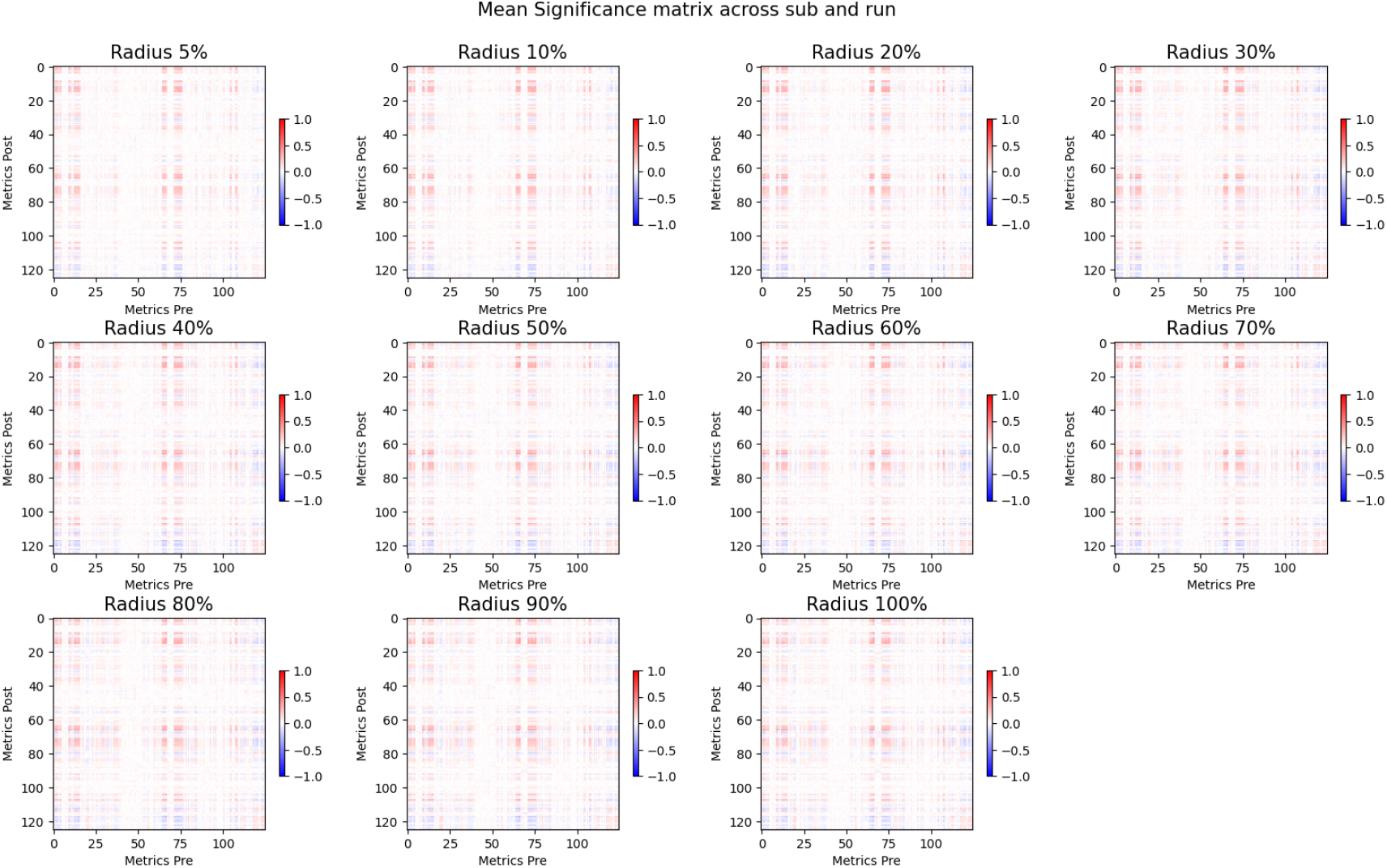
HDEEG data: Significance matrices for the full set of MOIs, averaged across all sessions, for varying radii. The matrices’ entries represent the fraction of sessions where the pre/post correlations were significant. Red (blue) represents values larger (smaller) than expected compared to the null model. Several (but not all) MOI pairs display significant levels of correlation between pre- and post-stimulus dynamics.

**Figure S5:**
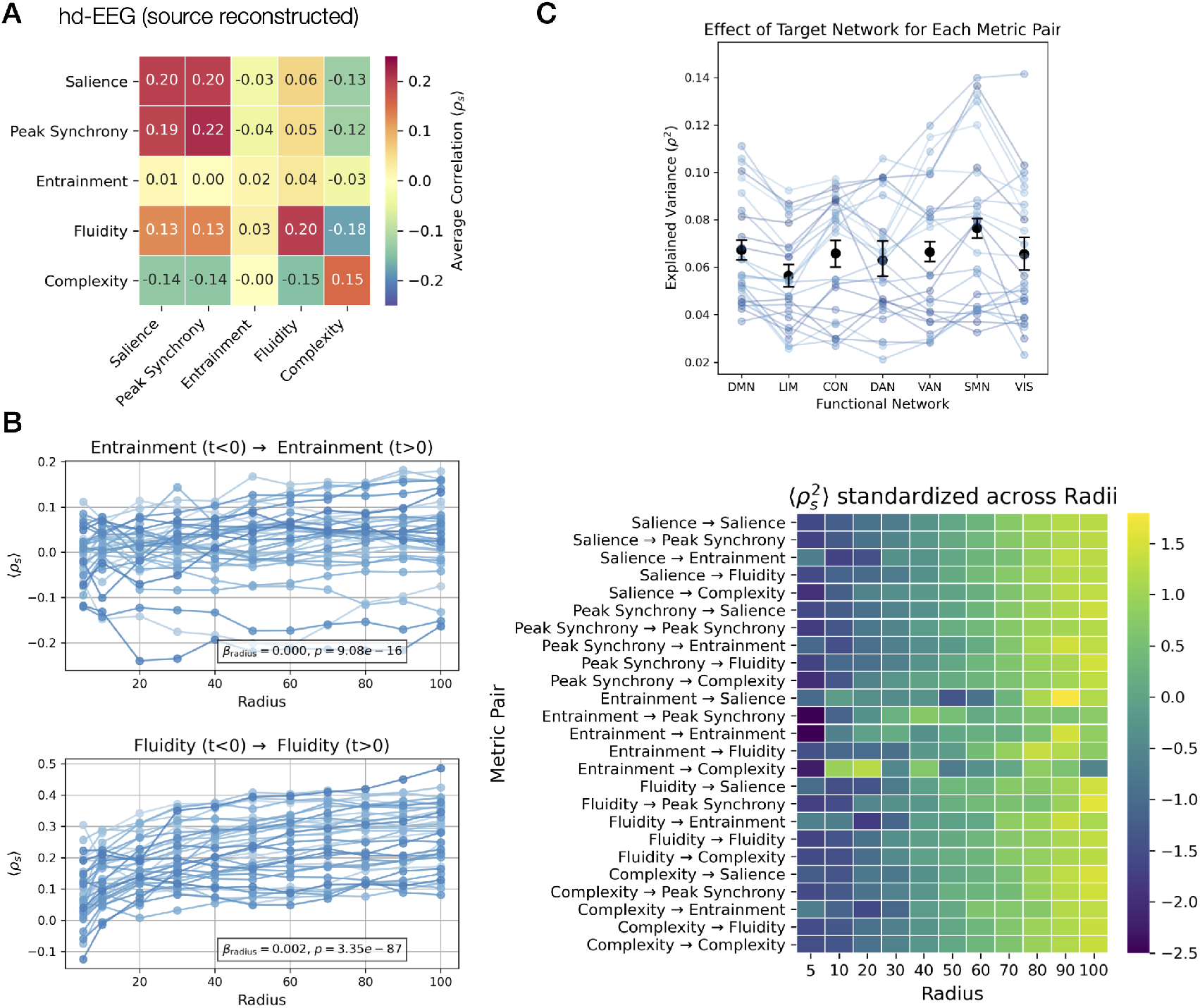
Main results reproduced for source reconstructed hd-EEG data. A) The average correlation between pre- and post-stimulus metrics (same as Fig.2). B) The dependence of the pre/post correlations as a function of the radius within which the pre-stimulus metrics were calculated (same as Fig.5). C) The dependence of the pre/post correlations as a function of the network stimulated (same as Fig.6).

**Figure S6:**
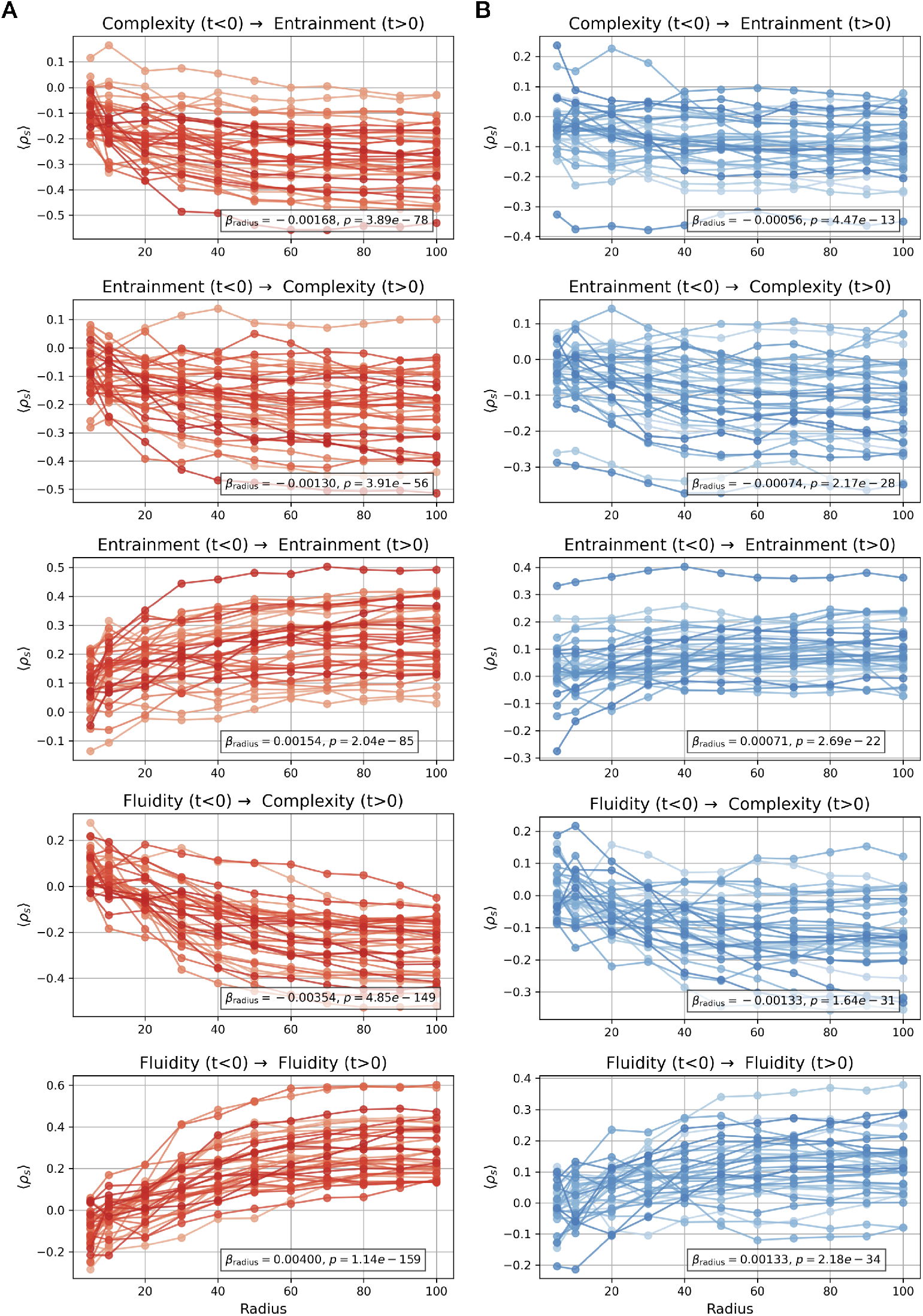
Example of radius dependence for a diverse pre/post metric pairs in SEEG modality (same as Fig.5). These demonstrate that the correlation between pre- and post-stimulus metrics 30 increases in absolute value as the radius increases.

**Figure S7:**
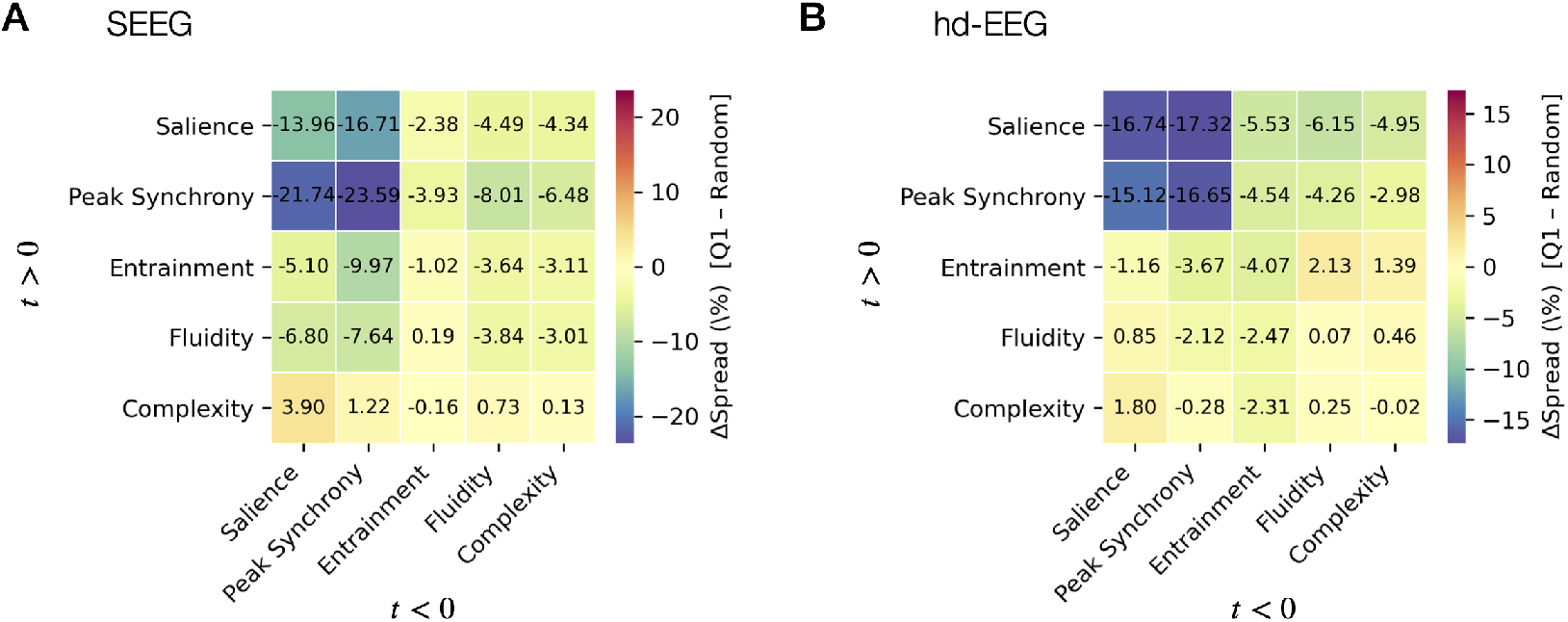
Heatmaps summarizing the mean Δ Spread across participants and sessions (as defined in Fig.4 for all pre–post metric pairs, in SEEG (A) and hd-EEG (B). Columns denote the conditioning pre-stimulus metric, rows the post-stimulus metric. Cooler colors indicate reduced variability when stimulation was conditioned on low pre-stimulus states.

